# Revealing spatio-temporal dynamics with long-term trypanosomatid live-cell imaging

**DOI:** 10.1101/2021.09.15.460474

**Authors:** Richard S. Muniz, Paul C. Campbell, Thomas E. Sladewski, Lars D. Renner, Christopher L. de Graffenried

**Affiliations:** Department of Molecular Microbiology and Immunology, Brown University, Providence, Rhode Island, USA; Leibniz Institute of Polymer Research and the Max Bergmann Center of Biomaterials, Dresden, Germany

**Author notes:** Address correspondence to Christopher de Graffenried. These authors contributed equally to this work (Richard S. Muniz, Paul C. Campbell).

## Abstract

*Trypanosoma brucei*, the causative agent of human African trypanosomiasis, employs a flagellum for dissemination within the parasite’s mammalian and insect hosts. *T. brucei* cells are highly motile in culture and must be able to move in all three dimensions for reliable cell division. These characteristics have made long-term microscopic imaging of live *T. brucei* cells challenging, which has limited our understanding of a variety of important cell-cycle events. To address this issue, we have devised an imaging approach that confines cells to small volumes that can be imaged continuously for up to 24 h. This system employs cast agarose microwells generated using a PDMS stamp that can be made with different dimensions to maximize cell viability and imaging quality. Using this approach, we have imaged individual *T. brucei* through multiple rounds of cell division with high spatial and temporal resolution. We have employed this method to study the differential rate of *T. brucei* daughter cell division and show that the approach is compatible with loss-of-function experiments such as small molecule inhibition and RNAi. We have also developed a strategy that employs in-well “sentinel” cells to monitor potential toxicity due to imaging. This live-cell imaging method will provide a novel avenue for studying a wide variety of cellular events in trypanosomatids that have previously been inaccessible.

**Importance:** *Trypanosoma brucei* causes severe diseases that affect humans and livestock in Sub-Saharan Africa. Efficient strategies for manipulating the *T. brucei* genome have provided a wealth of information about protein localization and function in diverse cellular processes. However, employing live-cell imaging for phenotypic analysis in *T. brucei* remains a significant challenge because immobilization of this highly motile parasite rapidly leads to morphologic defects and cell death. While fixed-cell imaging can provide snapshots of cellular events, it cannot provide the direct causal link or precise timing of events that comes from watching a living cell change over time. Our strategy using agarose microwells now allows long-term live cell *T. brucei* imaging with a simple apparatus that can be adapted for a wide range of experimental conditions.

## Introduction

Live-cell imaging has yielded vital information about protein function and the spatio-temporal dynamics of essential cellular events in a variety of organisms (1–3). Its key advantage is the ability to observe cells or structures within cells directly transition from one state to another, establishing a direct link between the initial, intermediate, and final states. This allows calculation of rates for many cellular processes, which is vital for understanding their function and how perturbations can alter their progression. While analysis of fixed samples can provide snapshots of individual events, live-cell imaging is essential for understanding dynamic cellular processes (4–6). Improvements in imaging hardware now enable live-cell imaging experiments that span from less than a second to many hours. However, long-term live-cell imaging of highly motile cells or cells that do not tolerate immobilization remains a challenge. Among the organisms that fall within these categories are the trypanosomatid parasites, which are important human pathogens that cause illnesses such as human African trypanosomiasis in Sub-Saharan Africa.

*Trypanosoma brucei* is an obligate extracellular parasite with an attached flagellum whose beat causes significant distortion of the entire cell body, which complicates live cell imaging (7, 8). Several approaches have been used for short-term imaging of live *T. brucei* cells, but these methods are not compatible with parasite division, producing apoptotic cells after less than 4 h (9–11). For long-term live-cell imaging, it is possible to gently immobilize the parasite on the surface of an agarose pad (12–14). However, cells imaged using this approach take much longer to divide than in conventional suspension culture and many arrest, suggesting that immobilization interferes with cellular processes. Long-term live-cell experiments, such as observing emerging RNAi phenotypes, have not been conducted due to the artefacts introduced by current imaging approaches.

One area that would benefit from live-cell imaging is the study of *T. brucei* cell division. The parasite maintains its highly polarized shape throughout the process, which leads to the ingression of a cleavage furrow along the long axis of the cell from the anterior to the posterior end. Cell division in *T. brucei* is intrinsically asymmetric and produces daughter cells that must remodel their cytoskeletons upon completion of cytokinesis prior to initiating another round of division (12). The daughter cell that inherits the new flagellum inherits the old-cell posterior and the new-cell anterior, while the old-flagellum daughter inherits the new-cell posterior and old-cell anterior. The new structures have distinct morphologies that are remodeled prior to subsequent rounds of cell division (13). The new flagellum is also shorter than the old flagellum at the completion of cell division, so it must be extended afterwards (14). It is possible that these remodeling events take different lengths of time, which could manifest as differing rates of cell division for the daughter cells.

There are many outstanding questions about the function of key cell-division regulators in *T. brucei* that could be addressed with long-term live-cell imaging. The Polo-like kinase homolog in *T. brucei* TbPLK is essential for the formation of a new flagellum attachment zone (FAZ), which attaches the flagellum to the cell body and directs placement of the cleavage furrow (15–17). In fixed samples, cells lacking TbPLK activity have severe cytokinetic defects and do not assemble a new FAZ, which leads to the production of looped and fully detached new flagella (18, 19). Due to a lack of live-cell imaging, it is not known if these two flagella phenotypes are produced by different mechanisms or if the looped flagella become fully detached over time. The cytokinetic protein TOEFAZ1 (also called CIF1) is a substrate and binding partner of TbPLK that localizes to the tip of the new FAZ and demarcates the location of cleavage furrow initiation (20, 21). TOEFAZ1 depletion causes severe cytokinetic defects. However, it has been suggested that in the absence of TOEFAZ1, cells can continue to divide using a “back-up” cytokinesis employing a furrow that ingresses from the posterior end (22). While we have observed remodeling of the parasite posterior end in cells lacking TOEFAZ1, our growth experiments strongly suggest that these cells are not able to complete viable cell divisions. Fixed-cell and short-term live cell imaging have shown some evidence of cells with posterior furrows, but entire cell divisions showing successful posterior-directed divisions have not been captured (22).

In this work, we have adapted a strategy from bacterial live-cell imaging that employs agarose microwells to confine *T. brucei* in small volumes without immobilization, which allows them to remain viable for up to 24 h while dividing at rates similar to those observed in bulk cultures. By observing the differential rates of daughter cell division, we show that the old-flagellum daughter cell produced by the *T. brucei* asymmetric cell division completes a subsequent round of cell division more rapidly than the other new-flagellum daughter. Our imaging approach is compatible with loss-of-function experiments such as small-molecule inhibition and RNAi, which we have employed to better understand the phenotypes that emerge when TbPLK and TOEFAZ1 are perturbed. We have also devised a strategy that allows us to include resistant “sentinel” cells within wells to serve as internal controls for imaging-associated toxicity. Overall, our imaging approach should be amenable for general use and will allow the study of many aspects of trypanosomatid biology that currently cannot be observed directly.

## Results

Agarose microwells have been used previously to isolate and image a variety of bacteria through multiple cell cycles (23, 24). Most of these approaches employ higher percentage agarose solutions to cast the microwell, followed by introduction of bacteria embedded in low-percentage agarose solutions to immobilize them for long-term imaging. Low-percentage agarose has been employed to immobilize the trypanosomatid *Leishmania* for live-cell imaging in the past, but using this strategy significantly impacts the speed and success rate of cell division events (25). We sought to develop a hybrid approach that uses agarose microwells to confine *T. brucei* cells in small volumes of liquid media, which should allow them to remain fully motile but unable to move outside of the imaging plane. The most important consideration is the size of the Z-dimension because dividing cells become significantly wider at late stages and need to be able to rotate to complete cytokinesis. However, providing too much space along this axis allows the parasite to move out of the focal plane, which impacts imaging.

The microwells are created from a polydimethylsiloxane (PDMS) stamp, which is generated from a photoresist master on a silicon wafer patterned with the wells using photolithography(26, 27). The designed pattern comprises many thousands of closely spaced wells, which is optimal for our cell plating strategy. Once the photoresist is exposed and developed on a silicon wafer substrate, PDMS is polymerized onto the pattern to produce its inverse, which then works as a master stamp when overlayed with agarose to recreate the original features present in the photoresist (**Figure S1A**). The PDMS stamp can be reused indefinitely. We generated a variety of PDMS stamps that create microwells ranging from 25 μm to 150 μm in the X and Y dimensions and 3 to 5 μm in the Z dimension to determine which sizes were compatible with long-term imaging.

We chose to focus on the imaging of an insect-resident form of *T. brucei*, known as the procyclic, because this form of the parasite does not require CO_2_ supplementation and tends to grow more easily in culture. To make wells suitable for *T. brucei*, we poured heated procyclic form (PCF) media supplemented with 10% FBS and 3.5% agarose onto the PDMS stamp and allowed it to cool (**Supplemental Figure S1B**). The excess agarose was cut away to produce a thin segment that fits within an imaging chamber slide. A 100 μL volume of media containing 1.6 million PCFs was then added to the bottom of the imaging chamber slide, followed by inversion of the gel slice onto the cells, trapping them in the wells (**Supplemental Figure S1C**). The slice is weighted down gently to keep the wells in tight contact with the surface of the imaging chamber, which keeps the cells from escaping from the wells. Once assembled, the chamber containing the microwells is overlaid with mineral oil to limit evaporation and then the imaging chamber slide is sealed with a glass slide and vacuum grease (**Supplemental Figure S1D-E**).

For imaging, we used an inverted microscope equipped with a motorized stage and a definite focus system to retain the imaging volume in focus for long periods of time. Since PCFs usually grow at slightly above room temperature without CO_2_ supplementation, we did not employ environmental control during imaging. The microscope is equipped with a sensitive sCMOS camera with a large sensor area (2048 × 2048 pixels), which allows a large field of view to be imaged. The camera is attached to a split-view system so that the camera sensor can simultaneously capture two image channels, which is necessary to register multichannel images due to the motility of the cells. The split-view can capture green and red fluorescence simultaneously or differential interference contrast (DIC) and a single fluorescent channel. We have used lenses ranging from 20× to 100× to image cells in microwells, depending on the precise experimental goals of the imaging. Our favored general approach uses a high numerical aperture (1.3 NA) oil-immersion 40× objective that provides an ideal mixture of resolution, light gathering, and large field of view.

As an initial test of the microwell imaging approach, we plated Lister-427 strain cells modified for tetracycline-inducible expression using the SmOx system (SmOx 427) in 100 × 100 × 5 μm wells and imaged them by DIC using the 40× lens, taking images every 10 min for 24 h (28). This well size allowed us to image 4 wells simultaneously with a 40× objective, of which 3 contained single cells. Over the course of 24 h, we observed multiple rounds of cell division in all the occupied wells, with minimal drifting of the microwells or cell death. The two neighboring wells that started with single cells are shown in **Figure 1A** (see **Movie S1** in the supplemental material). Over the course of 24 h, the cell number in the left well went from 2 to 9, while the lower well increased from 1 to 4. In conventional tissue culture, procyclic cells take approximately 8.5 h to undergo cell division, which would suggest that we were able to observe between two and three rounds of cell division, consistent with the expected cell cycle length. We have observed cells undergoing more than 3 rounds in a 24 h period. Since the single cells plated in each well were unlikely to be at the beginning of the cell cycle, it is likely that their first cell division took less than 8.5 h, which would account for what appeared to be elevated cell division rates. This suggests that DIC imaging and confinement in the wells does not inhibit cell proliferation over a 24 h time course.

**Figure 1.**
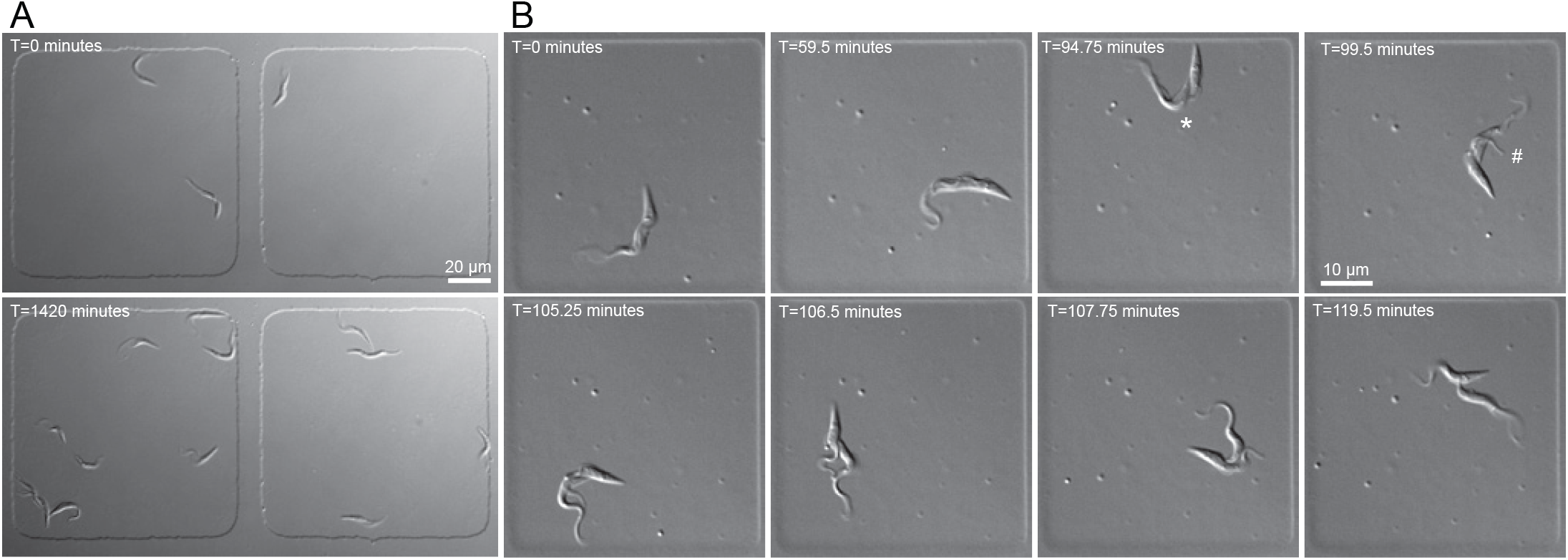
DIC imaging of procyclic *T. brucei* confined in agarose microwells. **A**. 1.6×10^6^ SmOx cells in 125 μL were plated and imaged over a 24 h time course in 100×100×5 μm wells. Cells were imaged every 10 min with a 40×/1.3 NA oil lens and 30 ms exposures. 3 initial cells result in 13 final cells after 24 h. **B**. Representative images of SmOx cells plated in 50×50×5 μm wells and imaged with a 40×/1.3 NA oil lens. DIC images were captured every 15 s for 2 h. * denotes division fold formation, # denotes cleavage furrow initiation.

Subsequent experiments showed that wells ranging from 50 to 150 μm in the X-Y dimensions provided robust cell divisions. We found that 5 μm in the Z dimension provided the best cell viability with cells occasionally drifting slightly out of focus for single exposures, while the 4 μm height provided better imaging with slightly fewer cell divisions. The 3 μm height chambers appeared to constrain the cells, which could improve short-term imaging runs but caused delays in cell division that makes them unsuitable for long-term imaging. We also found that the addition of 100 μM glutathione as a radical scavenger appeared to diminish imaging toxicity, especially when fluorescence imaging was employed (see below).

While we were able to capture many dividing cells in larger wells with a 10-min imaging increment, we wanted to determine if we could image single cells at higher sampling rates. To do this, we employed 50 × 50 × 4 μm wells and imaged wells every 15 s for 7.5 h (**Figure 1B** and **Movie S2**). Using these parameters, we were able to capture an entire *T. brucei* cell division cycle with high spatial and temporal resolution. This shows that DIC imaging is minimally toxic and that we can achieve sampling rates that are high enough to capture most cellular events.

With our DIC imaging strategy established, we sought to determine if the asymmetric daughter cells produced during *T. brucei* cell division subsequently divide at different rates. We imaged wells containing one SmOx 427 cell until it divided, then timed the length of each of the subsequent daughter cell divisions (**Figure 2A** and **Movie S3**). We noted that the rate of cell division for the two daughter cells was significantly different, with an average value of 91 min elapsing between the division of the first and the second daughter cells. The time required for 50% (T_1/2_) of the first daughter cells to divide is 510 min, whereas T_1/2_ for the second daughter cells to divide is 600 min (**Figure 2B**). This difference suggests that the two daughter cells divide at different rates. However, we were unable to distinguish between the new-flagellum daughter and old-flagellum daughter to determine if one of them persistently divided prior to the other, which would suggest that their remodeling took different lengths of time to reach the point where they begin to divide again.

**Figure 2.**
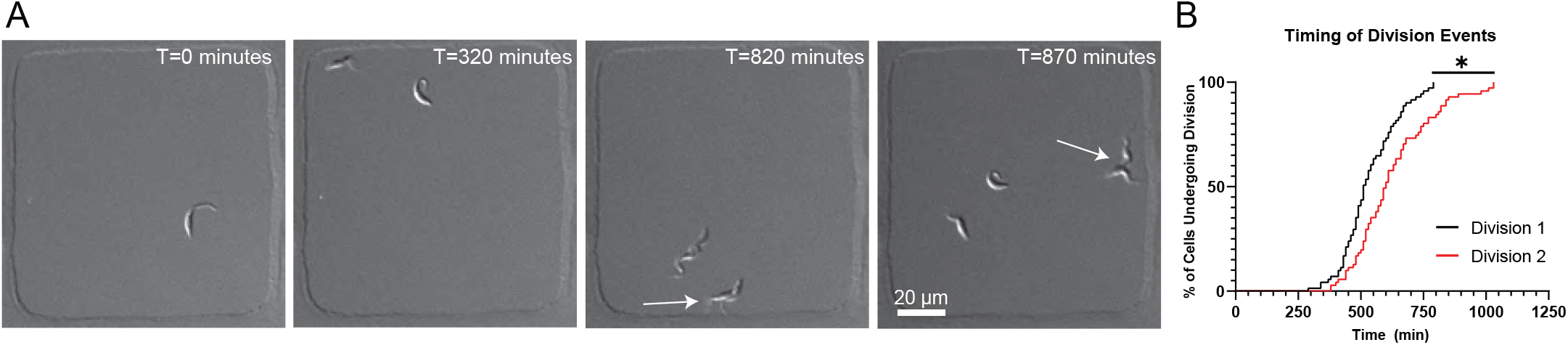
Daughter cells arising from an asymmetric division event have different timing to complete a subsequent cell cycle. **A**. Representative images from a 24 h time course of SmOX cells in 100×100×5 μm wells, imaged with a 20×/0.8 NA air lens. Initial cell divides at T=320 min. First daughter cell divides at T=820 min, followed by second daughter division at T=870 min (division events denoted by arrows). **B**. Staircase plot showing timing of first and second daughter cell divisions. T_1/2_ for Daughter Cell 1 division = 510 min, T_1/2_ for Daughter Cell 2 division = 600 min. Daughter 1 division average = 536 min, Daughter 2 division 2 average = 628 min. n=71 initial cells. *P<0.0001.

The most direct approach for distinguishing between the new-flagellum daughter and old-flagellum daughter cells would be to selectively mark the flagellum in a dividing cell. A previous approach to studying the division rate of the daughter cells depleted a non-essential flagellar axoneme protein using RNAi and modelled the persistence of a YFP-tagged allele of the depleted protein (13). We used a similar approach by C-terminally tagging one allele of the paraflagellar rod (PFR) protein PFR2 with the GFP variant mClover3 (mClv3) in SmOx 427 cells (29). The PFR is a paracrystalline structure closely associated with the flagellum that is thought to provide additional rigidity to the axoneme to facilitate *T. brucei* motility (30). Near-complete depletion of PFR2 causes severe motility defects in procyclic cells, but by targeting only the tagged allele with RNAi directed against mClv3, it should be possible to remove the PFR2-mClv3 from the cells selectively while leaving the wild-type (WT) pool of the protein intact (31, 32). In a dividing cell, this should leave the old flagellum mClv3 positive, while producing a non-fluorescent new flagellum (**Figure 3A**). We tested our depletion strategy in a cell line carrying a PFR2 allele C-terminally tagged with mClv3 and an inducible RNAi hairpin directed against mClv3. Control and mClv3 RNAi cells were fixed and labeled with the L8C4 monoclonal antibody against PFR2, which should detect both the WT and mClv3-tagged alleles (33). In the control samples, dividing cells had old and new flagella that were both mClv3 and L8C4 positive (**Figure S2B**). In cells where mClv3 RNAi had been induced for 12 h, we were able to observe dividing cells where both flagella were L8C4 positive, but only the old flagellum was mClv3 positive (**Figure S2B**). This showed that our specific depletion strategy differentially labels the old and new flagella. The growth of control and mClv3 RNAi cells was monitored in large-scale cultures over 48 h with cell counting. There was no appreciable change in growth between the samples and no evidence of cell sedimentation caused by motility defects, showing that the remaining pool of WT PFR2 was able to compensate for the absence of the mClv3-tagged allele (**Figure S2B**).

**Figure 3.**
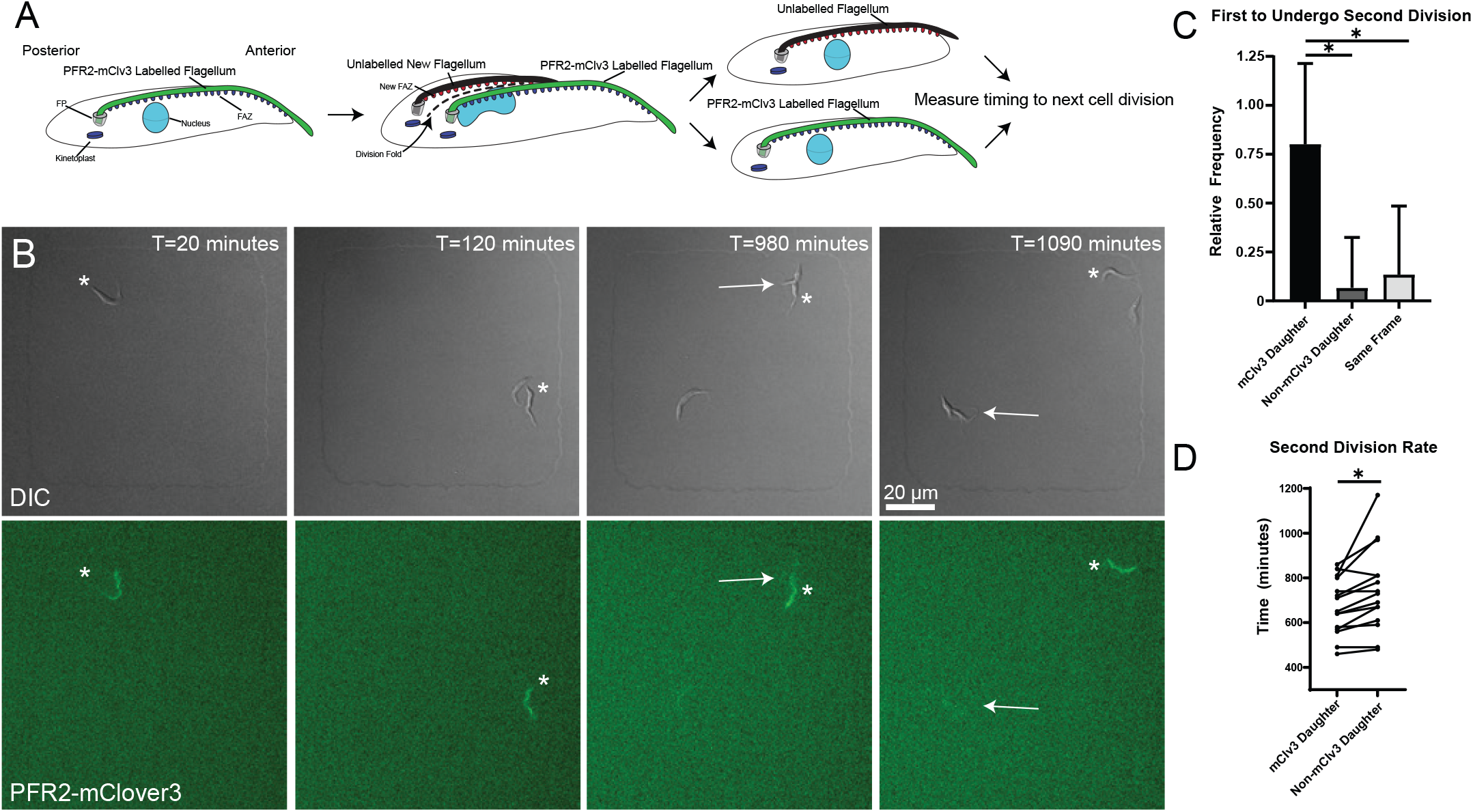
The old-flagellum daughter cell is able to complete a subsequent cell cycle before the new-flagellum daughter. **A**. Schematic depicting PFR2-mClv3 RNAi strategy. Induction of RNAi against PFR2-mClv3 results in an unlabelled new PFR. The first division event results in one daughter with a fluorescent PFR, and one non-fluorescent daughter. The two daughter cells are then tracked to determine which undergoes cytokinesis first. FP: Flagellar pocket. FAZ: Flagellum Attachment Zone **B**. PFR2-mClv3 RNAi was induced for 12 h before plating. Representative images from a 24 h time course of an initial fluorescent PFR2-mClv3 cell (denoted by * in all images). First division at T=120 min results in one cell with, and one without a fluorescent PFR structure. The fluorescent daughter cell undergoes a subsequent division (T=980 min) prior to the non-fluorescent daughter (T=1090 min). Arrows depict cytokinetic events. **C**. Relative frequencies of PFR2-mClv3 labelled and non-fluorescent daughter cells undergoing subsequent cell division first. 80% (n=12 cells) of instances the fluorescent PFR2-mClv3 daughter cell undergoes division first, 6.67% (n=1) non-fluorescent daughter cell divides first, and 13.3% (n=2) occur within the same frame. Error bars depict standard deviation. **D**. Direct comparison of timing of paired fluorescent and non-fluorescent daughter cells. Fluorescent daughter cells divided on average after 671 min, non-fluorescent daughter cells divided on average after 746 min. *P<0.0001.

To test if there is a difference in growth rate between the daughter cells, we induced mClv3 RNAi in our PFR2-mClv3 cell line for 12 h, which we found to be the point where the RNAi led to the production of unlabeled flagella. We plated the cells in microwell chambers and identified wells containing single cells with an mClv3-positive flagellum and a non-fluorescent flagellum that could be identified by DIC (**Figure 3B** and **Movies S4**). The unlabeled flagellum was always closer to the posterior of the dividing cell, which is where the new flagellum is located. We followed the cell employing simultaneous DIC and fluorescence microscopy as it completed division, which produced an mClv3-negative new-flagellum daughter and a mClv3-positive old-flagellum daughter, and then timed the division of the daughter cells. In 80% of the cases, the old-flagellum daughter cell underwent a subsequent round of division first, while in 13% of cases the two daughter cells divided within the same imaging interval of 10 min (**Figure 3C-D**). We observed only one case (6%) where the new-flagellum daughter divided first. The average time difference between the division of the two daughter cells was 75 min. These results suggest that there is a difference in the division rate of the asymmetric daughter cells produced during *T. brucei* cell division, with the process of remodeling the new cell posterior in the old-flagellum daughter cell occurring more quickly than remodeling the cell anterior and extending the flagellum to its complete length in the new-flagellum daughter cell.

One important goal for *T. brucei* live-cell imaging is the ability to observe phenotypes manifest during loss-of-function experiments. While we have attempted to make our live-cell imaging as minimally invasive as possible, it can be difficult to determine if there is some form of toxicity within a specific well that is contributing to an emerging phenotype. This toxicity could be due to illumination, which can cause cell damage by a variety of pathways, or due to diminishing nutrients in the media or the buildup of proteins or metabolic byproducts released by compromised cells. One way to address this issue would be to include cells in the imaging runs that are resistant to the drug or treatment being administered. These cells, which we will refer to as sentinels, would then serve as internal controls to monitor the conditions within the well. Any phenotypes that arise in both the sentinel and experimental cells would be attributable to the live-cell imaging and not the administered treatment. The sentinels should be easily distinguishable from the experimental cells so that phenotypes can be correctly assigned, which could be done by the expression of a cytosolic GFP to mark one cell population.

We have previously designed an analog-sensitive approach for the selective inhibition of the essential *T. brucei* kinase TbPLK (19). A mutation in the ATP binding site of the kinase produces a novel binding pocket that accommodates general kinase inhibitors modified with a bulky substituent, which generates orthogonal kinase-inhibitor pairs (34, 35). We previously showed that the bulky inhibitor 3MB-PP1 rapidly blocks cell division in analog-sensitive TbPLK (TbPLKas) cells but has no effect on wild-type 427 cells (19). We generated sentinel 427 cells expressing cytoplasmic mNeonGreen to serve as internal controls in imaging runs where TbPLKas activity was inhibited.

We treated TbPLKas and 427 sentinel cells with 2.5 μM 3MB-PP1 for 3 h, then plated them together at a 1:1 ratio in wells that contained the same concentration of drug (**Figure 4A-C** and **Movies S5**). The cells were imaged for 24 h at 10 min intervals using simultaneous DIC and fluorescence microscopy to distinguish between the TbPLKas and sentinel cells. TbPLK inhibition causes defects in the formation of a new flagellum attachment zone, which leads to detachment of the new flagellum and defects in cytokinesis (18). Over the course of 24 h, the 427 sentinel cells underwent multiple rounds of division, while the TbPLKas cells developed detached flagella and arrested, which is consistent with our previous results (19). We measured the growth of the cells by counting division events stemming from a single initial sentinel or TbPLKas cell at the beginning of the time course. Among the 427 sentinels, 88% of the cells underwent one round of cell division. We observed second division events 68% of the time and third events 60% of the time, while 20% underwent further rounds of division. For the TbPLKas cells, 29% completed one division. We observed second and third division events 11% and 3% of the time, respectively (**Figure 4B**). On average, 427 sentinel cells produced 2.36 division events, while the TbPLKas cells on average had 0.46 division events (**Figure 3C**). We also noted that TbPLKas cells initially produced looped flagella that became fully detached over time as the tip of the new flagellum released, which shows that the two previously established phenotypes arise from the same initial state (**Figure 4D** and **Movie S6**). This data shows that we can distinguish between phenotypes that arise from specific interventions, such as enzymatic inhibition by small molecules, and potential off-target effects caused by long-term imaging.

**Figure 4.**
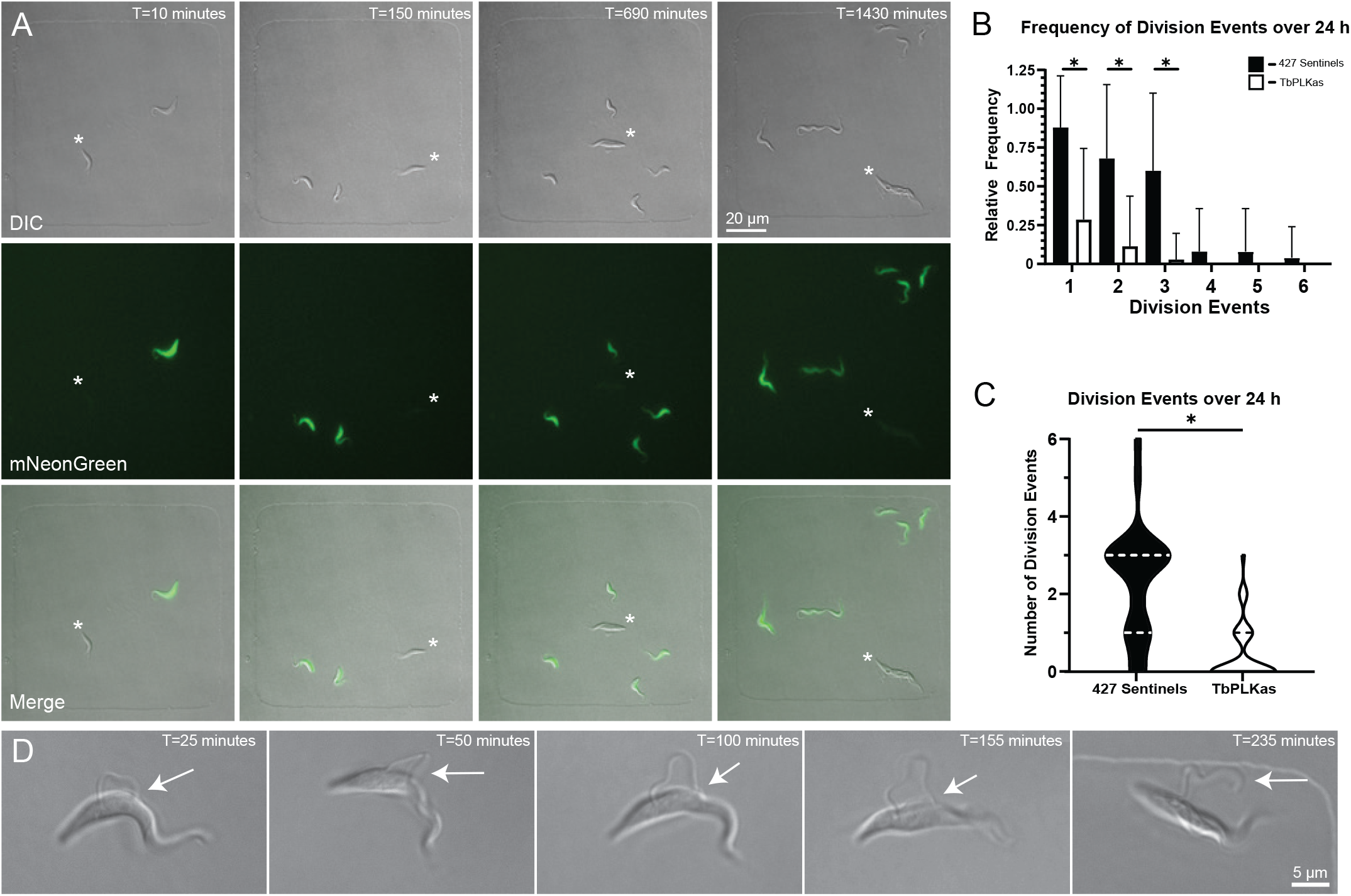
Small molecule inhibition of TbPLKas inhibits cytokinesis. TbPLKas and mNeonGreen expressing 427 sentinels were incubated with 2.5 μM 3MB-PPi for 3 h, followed by plating of 1×10^6^ cells from each condition in a total of 125 μL. Cells were imaged over 24 h, imaged every 10 min with 4% LED power, 30 ms exposure, and 2×2 pixel binning. **A**. Representative images from a 24 h time course of TbPLKas and 427 sentinel cells in 100×100×5 μm wells, imaged with a 40×/01.3 NA oil lens. Panels show a well containing one TbPLKas and one 427 sentinel cell. Over the course of 24 h, the 427-sentinel cell produces 5 total productive cell division events, while the TbPLKas cell (denoted in each panel by *) does not undergo a productive cell division. **B**. Relative frequencies of 427 sentinel (88% for D1, 68% D2, 60% D3, 8% D4, 8% D5, 4% D6 n=25) and TbPLKas (28.5% D1, 11.4% D2, 2.8% D3, 0% D4, 0% D5, 0% D6 n=35) cell division events 1 through 6 over 24 h. *P<0.0001. **C**. Truncated violin plot depicting average number of divisions for 427 sentinels (Average=2.360 divisions, 25% Percentile=1, 75% Percentile=6, n=25) and TbPLKas (Average=0.46 divisions, 25% Percentile=0, 75% Percentile=1, n=35). *P<0.0001. Error bars depict standard deviation. **D**. Insets depicting a looped-new flagellum phenotype in TbPLKas cells inhibited with 2.5 μM 3MB-PPi for 3 h prior to plating. 1×10^6^ cells were plated and imaged over 24 h, imaged every 5 min with 4% LED power, 30 ms exposure. Arrows point to tip of the new flagellum attached to the old flagellum from T=25 min to T=155 min. At T=235 min the new flagellum is completely detached along the length of the cell.

We previously showed that TOEFAZ1 RNAi caused a near-total block in cell division, with essentially all remaining cells showing aberrant morphologies and DNA content (20, 21). We did occasionally observe cells with what appeared to be remodeled posterior ends that still had attached anteriors that could be in the process of undergoing a posterior-initiated cytokinesis. To determine the progression of TOEFAZ1 RNAi in live cells, we induced RNAi against TOEFAZ1 in flasks for 16 h and then plated the cells in microwells to observe the emerging phenotypes (**Figure 5**). At this point in the RNAi time course the cells have begun to have division defects, but some cells still manage to ingress normal furrows from their anterior ends. We followed the division of multiple cells over the course of 20 h by DIC imaging. During our imaging, we observed 71 initial cells, of which 9 completed a single round of cell division. All the cells that managed to divide employed a furrow that had ingressed from the anterior end. We observed 3 cells that ingressed an anterior furrow that did not lead to a productive cell division during the length of imaging. We also identified 7 cells that appeared to remodel their posteriors in a manner similar to the reported posterior-initiated furrowing events (**Figure 5** and **Movie S7**) (22). However, none of these cells managed to complete cell division from the point where posterior furrowing was evident to the end of our imaging runs.

**Figure 5.**
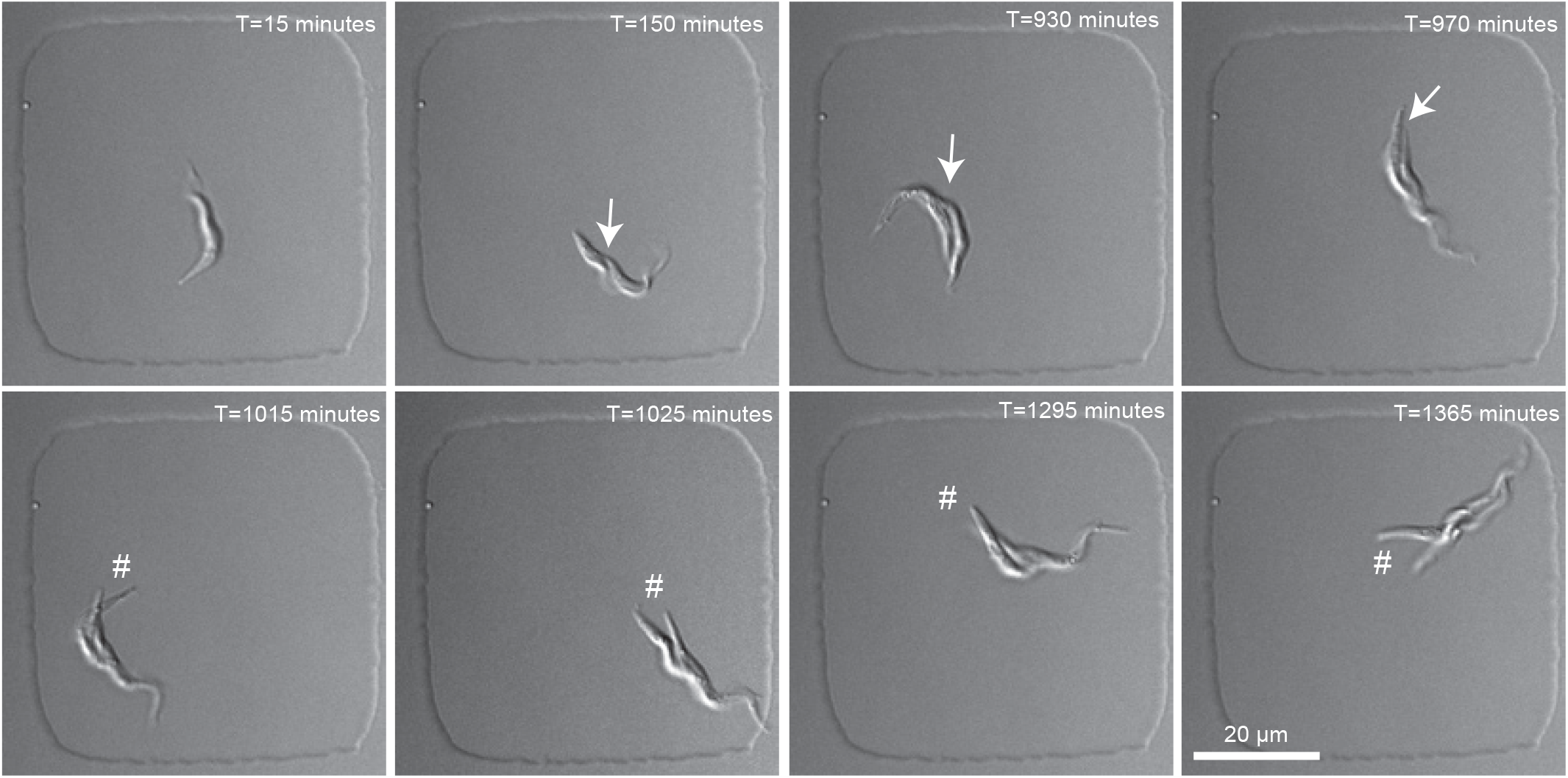
Cells depleted of TOEFAZ1 are not able to produce a productive cell division through a posterior furrow. TOEFAZ1-RNAi was induced for 16 h, after which 1.6×10^6^ cells in 125 μL were plated and imaged. **A**. A TOEFAZ1 depleted cell imaged over 24 h in a 50×50×5 μm well, with a 40×/1.3 NA oil lens imaged every 5 min with 30 ms exposure. A division fold is visible at T=150 min (denoted by arrow). Posterior furrow ingression is seen at T=1015 min (denoted by #) and persists for the remainder of the capture without a productive cell division.

As an additional control for our TOEFAZ1 RNAi runs, we generated SmOx cells expressing mClv3 under tetracycline-inducible control. These cells serve as sentinel cells for imaging experiments where a phenotype is triggered by tetracycline, such as the production of RNAi hairpins. We repeated our TOEFAZ1 RNAi imaging runs with our inducible SmOx sentinel cells expressing mClv3, following a 16 h induction of TOEFAZ1 RNAi and mClv3 sentinels (**Figure 6A-C** and **Movie S8**). We measured the growth of the cells by counting division events stemming from a single initial sentinel or TOEFAZ1 cell at the beginning of the time course. Among the SmOx mClv3-sentinels, 94% of the cells underwent cell division. We observed second division events 69% of the time and third division events 50% of the time, while 12.5% underwent subsequent division events over 20 h. For the TOEFAZ1 RNAi cells, only 30% completed one division event, and no cells were able to undergo subsequent cell divisions over the same period (**Figure 6B**). Over 20 h, induced SmOx sentinel cells averaged 2.3 division events, while TOEFAZ1 RNAi cells averaged 0.3 division events (**Figure 4C**). Importantly, we found that SmOx sentinel cells take at most 40 min (average 20 min) to complete furrow ingression following formation of the division fold, which is a structure that becomes visible shortly before the initiation of cytokinesis and defines the plane of furrow ingression (36). In contrast, the subset of TOEFAZ1 RNAi cells that formed posterior-initiated furrows generated division folds that persisted for an average 615 min (range 235-925 min). The cells with posterior furrow ingression (n=7) were unable to divide by the end of the imaging run, which averaged an additional 495 min (range 160-1065 min) from posterior furrow initiation. In cells that formed a posterior-initiated furrow, the shortest time between division fold formation and completion of an imaging run without a productive division event was 835 min, which contrasts with the maximum of 40 min we observed in SmOx sentinel cells. This strongly suggests that posterior-initiated furrows cannot produce viable cell divisions and do not represent an alternate pathway for cytokinesis in *T. brucei*.

**Figure 6.**
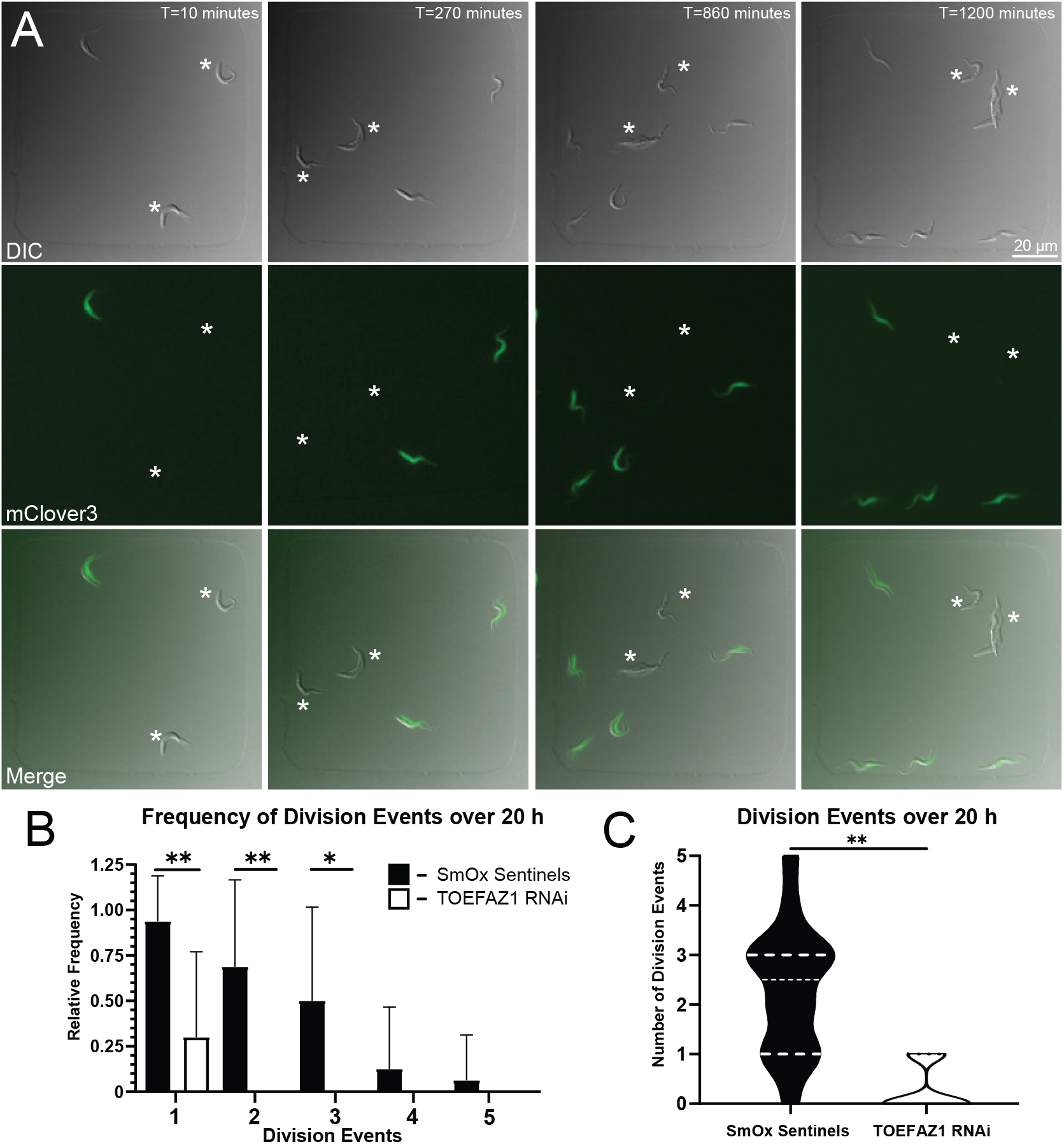
Depletion of TOEFAZ1 results in a rapid cessation of division. TOEFAZ1-RNAi and inducible SmOx mClv3 sentinel cells were induced for 12 h, after which 1×10^6^ cells from each condition were plated and imaged over 20 h in 100×100×5 μm wells with a 40×/1.3 NA oil lens and imaged every 10 min with 30% LED power and 30 ms exposure. **A**. Representative frames portrays 1 SmOx sentinel cell producing 3 productive division events, while 2 TOEFAZ1-RNAi cells (denoted by *) resulted in no productive division events. **B**. Relative frequencies of SmOx sentinels (93.8% D1, 68.8% D2, 50% D3, 12.5% D4, 6.3% D5, n=16) and TOEFAZ1-RNAi (30% for D1, 0% D2, 0% D3, 0% D4, 0% D5, n=20) cell division events 1 through 5 over 20 h. **P<0.0001, *P=0.0001. Error bars depict standard deviation. **C**. Truncated violin plot depicting average number of division events for SmOx sentinels (Average=2.3 divisions, 25% Percentile=1, 75% Percentile=3, n=16) and TOEFAZ1-RNAi (Average=0.3 divisions, 25% Percentile=0, 75% Percentile=1, n=20). **P<0.0001.

## Discussion

In this work, we have devised a strategy using agarose microwells that addresses a long-standing need in *T. brucei* phenotypic analysis: the ability to image live cells continuously for up to 24 h. Our approach is compatible with both DIC and fluorescence imaging, can vary the degree of parasite confinement for different imaging needs, and is sufficiently mild to allow the assessment of phenotypes that emerge during loss-of-function experiments. The degree of in-well toxicity can be measured using sentinel cells, which provides additional confidence for phenotypic analysis. We have used our live-cell imaging approach to address several questions pertaining to *T. brucei* cell division that have not been feasible previously. We show that the old-flagellum daughter cell appears to divide more quickly than the new-flagellum daughter, that the fully detached flagella observed upon TbPLK inhibition are initially produced as looped flagella, and that the posterior “furrowing” events that occur in cells lacking TOEFAZ1 do not appear to lead to cell division events. These results show the efficacy of our imaging approach and how it can be employed to study novel aspects of *T. brucei* biology.

The materials necessary for our live-cell imaging method should be accessible to many labs. The design of the microchambers on a silicon wafer substrate can be generated using CAD files, readily replicated into PDMS stamps and customized for specific experimental needs. Beyond media and agarose, the imaging chamber slides are the only other significant consumable expense. For imaging, a definite focus for maintaining the imaging plane is extremely helpful and is a common feature of most live-cell imaging systems. Capturing a single fluorescent channel or DIC imaging is straightforward, but fairly short (≤ 30 msec) exposures are necessary to limit blurring due to cell motility. This is not an issue for DIC imaging, where the power of the white-light source can be increased without any deleterious effects on the cells; bright fluorescent signals from endogenously tagged proteins or overexpression constructs are readily visible as well. Dual-channel imaging is more challenging because both channels must be captured simultaneously due to cell motility. A split-view system can solve this issue using a single camera, while newer microscopes can come equipped with multiple cameras and optimized light paths for simultaneous capture of multiple signals.

We showed that the old-flagellum daughter cell produced in the asymmetric *T. brucei* cell division appears to divide more rapidly than the new-flagellum daughter cell. This suggests that the extension of the new flagellum and remodeling of the new cell anterior require more time to accomplish than remodeling of the new cell posterior, which is inherited by the old-flagellum daughter cell. This is consistent with the degree of cellular remodeling that must be performed by the two daughter cells. Reshaping the ends of the cell body requires adjustments to the subpellicular microtubule array, which underlies the plasma membrane and is responsible for *T. brucei* morphology (13). These adjustments likely involve the addition of new microtubules to the array or extension of extant microtubules. The new anterior end must be extended to diminish the portion of the flagellum that overhangs the cell body. This remodeling likely requires extension of the FAZ to attach the flagellum to the cell body, as does the lengthening of the new flagellum. The remodeling of the posterior end does not require any adjustments to the flagellum or FAZ, which likely explains why the old-flagellum daughter is able to re-enter cell division slightly faster than the new-flagellum daughter. Any consequences of this growth difference are currently unknown, although it should be noted that there are several distinct asymmetric cell division events that occur as the parasite changes life cycle stages as it transits through the insect vector (37, 38). Differences in the division rates of the daughter cells produced by these uniquely configured cell divisions may skew the population towards one life cycle stage.

TbPLK inhibition using the analog-sensitive approach produces parasites with flagellar detachment defects due to the absence of the new FAZ. The new and old flagella are linked by the flagella connector (FC), which is a multidomain structure found at the tip of the new flagellum that tracks along the length of the old flagellum as the new flagellum extends during cell division (39, 40). The FC disengages shortly after the initiation of cleavage furrow ingression to release the new flagellum (36). Unlike the FAZ, FC assembly does not appear to rely on TbPLK activity, so it is likely that the looped flagella we observe at early stages of cell division are linked by only the FC (20). At later stages of cell division, it is likely that the force of the beating flagellum is sufficient to disrupt the FC, which leads to new flagellum detachment. It is also possible that the FC linkage resolves normally once the new flagellum has reached a certain length. However, we frequently saw cells with fully detached flagella that had not formed a cleavage furrow division fold, so it appears that the forced rupture of the FC is more likely to occur than its normal release.

It has been proposed that cells lacking TOEFAZ1 can undergo cell division using a “back-up” approach that generates a cleavage furrow that initiates at the cell posterior and then moves towards the anterior end (22). While backup cytokinetic mechanisms do occur in some organisms, they tend to have limited cell polarity and have the capacity to adhere to surfaces, which allows these organisms to use a pulling force to segregate the cellular organelles (41–44). In our live-cell experiments, we observed several TOEFAZ1 RNAi cells that appeared to form what had previously been described as posterior furrows. These cells were not able to complete furrow ingression, which occurred in control sentinel cells on average within 20 min (max 40 min), over the course of many hours. This strongly argues that the posterior furrows do not represent an alternate cytokinetic mechanism and instead are a “dead-end” phenotype. The remodeling of the cell posterior to produce a new posterior end is a distinct event from the ingression of the cleavage furrow, which makes it possible that posterior remodeling is proceeding in TOEFAZ1 RNAi cells in the absence of furrow formation (36). Considering the high degree of spatial and temporal organization in *T. brucei* cell division, it is difficult to envision how posterior furrow ingression would proceed and provide a viable cell division. In previous work, cells that form posterior furrows tended to have elevated DNA content, which suggests that the parasites have not successfully completed cell division and instead had restarted the process from G1 (22). This again suggests that posterior furrowing does not provide a viable pathway for the production of daughter cells.

We have shown that our live-cell approach can be applied to a wide range of potential experimental setups. Imaging the bloodstream form of the parasite will be an important future advance, along with the development of a DNA marker that allows us to monitor cell cycle progression with more precision. Future experiments will include monitoring parasites as they undergo life-cycle transitions to determine if these transformations require cell divisions to occur, which is an open question. The mechanisms that anti-trypanocidal drugs employ to trigger cell death could be directly observed using our approach. Imaging the related trypanosomatids *Trypanosoma cruzi* and *Leishmania* should be feasible and would allow comparative studies with *T. brucei* along with exploring the unique aspects of their biology. Another potentially exciting avenue would employ recently developed methods for simultaneous high-speed imaging of trypanosomatids on multiple Z-planes (45). Low frequency imaging by fluorescence or DIC could be used to wait for specific events to occur, such as specific stages of cell division, which could then be captured with higher sampling rates. Strategies for adding or removing material from individual wells would also open up new experimental avenues.

## Materials and Methods

Detailed descriptions of the antibodies employed, cell culture conditions, immunofluorescence, RNAi, statistical analysis, and microfabrication of the PDMS stamps are included in the supplemental materials.

### Microscopy

Images were taken using a Zeiss Axio Observer.Z1 microscope (Carl Zeiss Microscopy, Oberkochen, Germany) equipped with 20×/0.8 NA, 40×/1.3 NA, 63×/1.4 NA and 100×/1.4 NA Plan Aprochromat objective lenses. A W-View Gemini optical image splitter (Hamamatsu Photonics) was used for simultaneous dual imaging of differential interference contrast (DIC) and mClv3/mNeonGreen. Cells were simultaneously illuminated with 624/40 filtered transmitted light and excited with an X-Cite 120 LED illumination source (Excelitas Technologies, Waltham, MA) filtered with a 520/35 bandpass filter. Transmitted and excited light were then passed through a GFP/mCherry 520/35, 624/40 dual band cube filter and separated in the W-View Gemini optical image splitter with a 562 long pass dichroic filter. mNeonGreen/mClv3 wavelengths were then filtered through a 520/35 bandpass filter and the mCherry wavelengths were passed through a polarized filter to produce the DIC image. A Definite Focus.2 (Carl Zeiss Microscopy, Oberkochen, Germany) system was used to compensate for focus drift over the course of imaging. Images were obtained on an ORCA-Flash 4.0 V2 sCMOS camera (Hamamatsu, Shizuoka, Japan) running SlideBook 6 digital microscopy software (Intelligent Imaging Innovations, Inc.).

Immunofluorescence images were taken using a Plan Aprochromat 100x/1.4 NA oil lens. All images were analyzed with ImageJ (National Institutes of Health, Bethesda, MD), and assembled for publication in Adobe Photoshop and Illustrator (CC 2021).

### Generating agarose microwells

An 8% SeaPlaque GTG Agarose (Lonza, Basel, Switzerland) solution in Ultra Pure Water (Genesee Scientific, El Cajon, CA) was placed in an autoclave to ensure even heating. Simultaneously, base media for either 427- or SmOx-based cell lines was warmed to 37 °C. Agarose was added to the heated media to a final concentration of 3.5% agarose. The solution was briefly vortexed to mix, pulsed in a centrifuge to remove any bubbles, then pipetted onto the PDMS stamp containing impressions for desired well sizes and was allowed to cool. Agarose grids were inverted on a sterile surface and cut to size. Agarose grids that are not used day of can be stored at 4 °C with enough supplemental base media to prevent agarose grid drying out, for up to a week.

### SmOx live cell plating and imaging

SmOx cells were harvested by centrifugation at 800× g for 10 min at RT. The cells were resuspended in 1 mL base SmOx media with 100 μM of reduced L-glutathione (Sigma-Aldrich, St. Louis, MO). A 125 μL volume of cells containing 1 x10^6^ were plated in Lab-Tek II chambered coverglass #1.5 Borosilicate (Nunc International, Rochester, NY). The trimmed down agarose grid was inverted onto cells and gently weighed down to ensure consistent contact with base of chamber. Mineral oil was added to prevent evaporation and enclosed with a 24×60mm No1. Gold Seal Cover Slip (Thermo Scientific, Portsmouth, NY) sealed with high vacuum grease (Dow Corning, Midland, MI). For long term DIC imaging, cells were imaged every 5 or 10 min for 24 h with 30 ms exposures. For shorter duration, rapid DIC acquisition, cells were imaged every 15 s for 7.5 h with 30 ms exposures.

### Small drug inhibition of TbPLKas

Cultures of TbPLKas and mNeonGreen expressing 427 Sentinels in log-phase growth were seeded separately at 3×10^6^ cells/mL. 2.5 μM 3MB-PPi (APExBIO, Houston, Tx) dissolved in DMSO was added to the cultures for 3 h at 27° C before plating.

### Dual-channel imaging

For PFR2-mClv3, mClv3 RNAi, 1×10^6^ induced cells were plated in a 125 μL volume. Cells were identified with an existing fluorescent paraflagellar rod and imaged over 24 h at RT using a 40×/1.3 NA oil objective, imaging every 10 min with simultaneous split-DIC/Fluorescent imaging at 4% LED power, 30 ms exposure, and 2×2 pixel binning.

For TOEFAZ1 RNAi cells grown with SmOx sentinels, 1×10^6^ cells of both inducible pLew100-mClv3 SmOx sentinels and TOEFAZ1-RNAi cells were plated in a 125 μL volume. Cells were imaged over 20 h, using 20×/0.8 NA air or a 40×/1.3 NA oil objective, imaging every 5 or 10 min at RT with simultaneous split-DIC/Fluorescent imaging at 30% LED power, with 30 ms exposures.

For TbPLKas inhibition with 3MB-PPi, 8×10^6^ cells of both PLKas and 427Sentinels were mixed and harvested by centrifugation at 800 × g for 10 min at RT. Cells were resuspended in 1 mL of base 427 media with 2.5 μM 3MB-PPi, and 100 μM reduced L-glutathione. 125 μL of cell mixture were plated as previously described. Cells were imaged over 24 h, using 40×/1.3 NA oil objective, imaging every 10 min with simultaneous split-DIC/Fluorescent imaging at 4% LED power, with 30 ms exposures and 2×2 pixel binning.

## Acknowledgements

We would like to thank Professor Sue Vaughan and Dr. Jack Sunter for their input. The research reported in this publication was supported by NIAID-NIH under awards R21AI151490 and R01AI112953 to CLdG. The content is solely the responsibility of the authors and does not necessarily represent the official views of the National Institutes of Health.

## REFERENCES

1. Ettinger A, Wittmann T. 2014. Fluorescence live cell imaging, 1st ed. Elsevier Inc.

2. Bolbat A, Schultz C. 2016. Recent developments of genetically encoded optical sensors for cell biology. Biol Cell 109:1 23.

3. Specht EA, Braselmann E, Palmer AE. 2017. A Critical and Comparative Review of Fluorescent Tools for Live-Cell Imaging. Annu Rev Physiol 79:93 117.

4. Zoncu R, Perera RM, Sebastian R, Nakatsu F, Chen H, Balla T, Ayala G, Toomre D, Camilli PVD. 2007. Loss of endocytic clathrin-coated pits upon acute depletion of phosphatidylinositol 4,5-bisphosphate. Proc National Acad Sci 104:3793 3798.

5. Rosendale M, Perrais D. 2017. International Journal of Biochemistry and Cell Biology. Int J Biochem Cell Biology 93:41 45.

6. Lim TC, Hatano T, Kamnev A, Balasubramanian MK, Chew TG. 2018. Equatorial Assembly of the Cell-Division Actomyosin Ring in the Absence of Cytokinetic Spatial Cues. Curr Biol 28:955 962.e3.

7. Weiße S, Heddergott N, Heydt M, Pflästerer D, Maier T, Haraszti T, Grunze M, Engstler M, Rosenhahn A. 2012. A quantitative 3D motility analysis of Trypanosoma brucei by use of digital in-line holographic microscopy. Plos One 7:e37296.

8. Heddergott N, Krüger T, Babu SB, Wei A, Stellamanns E, Uppaluri S, Pfohl T, Stark H, Engstler M. 2012. Trypanosome Motion Represents an Adaptation to the Crowded Environment of the Vertebrate Bloodstream. PLoS pathogens 8:e1003023.

9. Hochstetter A, Stellamanns E, Deshpande S, Uppaluri S, Engstler M, Pfohl T. 2015. Microfluidics-based single cell analysis reveals drug-dependent motility changes in trypanosomes. Lab Chip 15:1961 1968.

10. Voyton CM, Choi J, Qiu Y, Morris MT, Ackroyd PC, Morris JC, Christensen KA. 2019. A Microfluidic-Based Microscopy Platform for Continuous Interrogation of Trypanosoma brucei during Environmental Perturbation. Biochemistry-us 58:875 882.

11. Price HP, MacLean L, Marrison J, O’Toole PJ, Smith DF. 2010. Validation of a new method for immobilising kinetoplastid parasites for live cell imaging. Mol Biochem Parasit 169:66 69.

12. Wheeler RJ, Gull K, Sunter JD. 2019. Coordination of the Cell Cycle in Trypanosomes. Annu Rev Microbiol 73:133–154.

13. Abeywickrema M, Vachova H, Farr H, Mohr T, Wheeler RJ, Lai D, Vaughan S, Gull K, Sunter JD, Varga V. 2019. Non-equivalence in old- and new-flagellum daughter cells of a proliferative division in Trypanosoma brucei. Mol Microbiol https://doi.org/10.1111/mmi.14345.

14. Farr H, Gull K. 2009. Functional studies of an evolutionarily conserved, cytochrome b5 domain protein reveal a specific role in axonemal organisation and the general phenomenon of post-division axonemal growth in trypanosomes. Cell Motil Cytoskel 66:24 35.

15. Kumar P, Wang CC. 2006. Dissociation of cytokinesis initiation from mitotic control in a eukaryote. Eukaryot Cell 5:92 102.

16. Hammarton TC, Kramer S, Tetley L, Boshart M, Mottram JC. 2007. Trypanosoma brucei Polo-like kinase is essential for basal body duplication, kDNA segregation and cytokinesis. Mol Microbiol 65:1229 1248.

17. Graffenried CL de, Ho HH, Warren G. 2008. Polo-like kinase is required for Golgi and bilobe biogenesis in Trypanosoma brucei. J Cell Biology 181:431 438.

18. Ikeda KN, Graffenried CL de. 2012. Polo-like kinase is necessary for flagellum inheritance in Trypanosoma brucei. J Cell Sci 125:3173 3184.

19. Lozano-Núñez A, Ikeda KN, Sauer T, Graffenried CL de. 2013. An analogue-sensitive approach identifies basal body rotation and flagellum attachment zone elongation as key functions of PLK in Trypanosoma brucei. Mol Biol Cell 24:1321 1333.

20. McAllaster MR, Ikeda KN, Lozano-Núñez A, Anrather D, Unterwurzacher V, Gossenreiter T, Perry JA, Crickley R, Mercadante CJ, Vaughan S, Graffenried CL de. 2015. Proteomic identification of novel cytoskeletal proteins associated with TbPLK, an essential regulator of cell morphogenesis in Trypanosoma brucei. Mol Biol Cell 26:3013 3029.

21. Sinclair-Davis AN, McAllaster MR, Graffenried CL de. 2017. A functional analysis of TOEFAZ1 uncovers protein domains essential for cytokinesis in Trypanosoma brucei. J Cell Sci 130:3918 3932.

22. Zhou Q, Gu J, Lun Z-R, Ayala FJ, Li Z. 2016. Two distinct cytokinesis pathways drive trypanosome cell division initiation from opposite cell ends. Proc National Acad Sci 113:3287 3292.

23. Tuson HH, Auer GK, Renner LD, Hasebe M, Tropini C, Salick M, Crone WC, Gopinathan A, Huang KC, Weibel DB. 2012. Measuring the stiffness of bacterial cells from growth rates in hydrogels of tunable elasticity. Mol Microbiol 84:874 891.

24. Eun Y-J, Ho P-Y, Kim M, LaRussa S, Robert L, Renner LD, Schmid A, Garner E, Amir A. 2018. Archaeal cells share common size control with bacteria despite noisier growth and division. Nat Microbiol 3:1 9.

25. Wheeler RJ, Gluenz E, Gull K. 2011. The cell cycle of Leishmania: morphogenetic events and their implications for parasite biology. Mol Microbiol 79:647 662.

26. Weibel DB, DiLuzio WR, Whitesides GM. 2007. Microfabrication meets microbiology. Nat Rev Microbiol 5:209 218.

27. Xia Y, Whitesides GM. 1998. Soft Lithography. Angewandte Chemie Int Ed 37:550–575.

28. Poon SK, Peacock L, Gibson W, Gull K, Kelly S. 2012. A modular and optimized single marker system for generating Trypanosoma brucei cell lines expressing T7 RNA polymerase and the tetracycline repressor. Mol Biochem Parasit 2:110037 110037.

29. Bajar BT, Wang ES, Lam AJ, Kim BB, Jacobs CL, Howe ES, Davidson MW, Lin MZ, Chu J. 2016. Improving brightness and photostability of green and red fluorescent proteins for live cell imaging and FRET reporting. Sci Rep-uk 6:1 12.

30. Alves AA, Gabriel HB, Bezerra MJR, Souza W de, Vaughan S, Cunha-e-Silva NL, Sunter JD. 2020. Control of assembly of extra-axonemal structures: the paraflagellar rod of trypanosomes. J Cell Sci 133:jcs242271.

31. Bastin P, Sherwin T, Gull K. 1998. Paraflagellar rod is vital for trypanosome motility. Nature 391:548–548.

32. Bastin P, Pullen TJ, Sherwin T, Gull K. 1999. Protein transport and flagellum assembly dynamics revealed by analysis of the paralysed trypanosome mutant snl-1. Journal of cell science 112 (Pt 21):3769 3777.

33. Kohl L, Sherwin T, Gull K. 1999. Assembly of the paraflagellar rod and the flagellum attachment zone complex during the Trypanosoma brucei cell cycle. J Eukaryot Microbiol 46:105 109.

34. Bishop AC, Ubersax JA, Petsch DT, Matheos DP, Gray NS, Blethrow J, Shimizu E, Tsien JZ, Schultz PG, Rose MD, Wood JL, Morgan DO, Shokat KM. 2000. A chemical switch for inhibitor-sensitive alleles of any protein kinase. Nature 407:395 401.

35. Zhang C, Kenski DM, Paulson JL, Bonshtien A, Sessa G, Cross JV, Templeton DJ, Shokat KM. 2005. A second-site suppressor strategy for chemical genetic analysis of diverse protein kinases. Nat Methods 2:435 441.

36. Wheeler RJ, Scheumann N, Wickstead B, Gull K, Vaughan S. 2013. Cytokinesis in Trypanosoma brucei differs between bloodstream and tsetse trypomastigote forms: implications for microtubule-based morphogenesis and mutant analysis. Mol Microbiol 90:1339 1355.

37. Sharma R, Peacock L, Gluenz E, Gull K, Gibson W, Carrington M. 2008. Asymmetric cell division as a route to reduction in cell length and change in cell morphology in trypanosomes. Protist 159:137 151.

38. Rotureau B, Subota I, Buisson J, Bastin P. 2012. A new asymmetric division contributes to the continuous production of infective trypanosomes in the tsetse fly. Development 139:1842 1850.

39. Moreira-Leite FF, Sherwin T, Kohl L, Gull K. 2001. A trypanosome structure involved in transmitting cytoplasmic information during cell division. Science 294:610 612.

40. Briggs LJ, McKean PG, Baines A, Moreira-Leite F, Davidge J, Vaughan S, Gull K. 2004. The flagella connector of Trypanosoma brucei: an unusual mobile transmembrane junction. J Cell Sci 117:1641 1651.

41. Choudhary A, Lera RF, Martowicz ML. 2013. Interphase cytofission maintains genomic integrity of human cells after failed cytokinesis.

42. Gerisch G, Weber I. 2000. Cytokinesis without myosin II. Curr Opin Cell Biol 12:126 132.

43. Rancati G, Pavelka N, Fleharty B, Noll A, Trimble R, Walton K, Perera A, Staehling-Hampton K, Seidel CW, Li R. 2008. Aneuploidy Underlies Rapid Adaptive Evolution of Yeast Cells Deprived of a Conserved Cytokinesis Motor. Cell 135:879 893.

44. Uyeda TQ, Nagasaki A. 2004. Variations on a theme: the many modes of cytokinesis. Curr Opin Cell Biol 16:55 60.

45. Walker BJ, Wheeler RJ. 2019. High-speed multifocal plane fluorescence microscopy for three-dimensional visualisation of beating flagella. J Cell Sci 132:jcs.231795.

