## Supplemental material for "Revealing spatio-temporal dynamics with long-term trypanosomatid live-cell imaging"

**Antibodies**

The antibody L8C4 from Keith Gull (University of Oxford- Oxford, United Kingdom) was used at a 1:100 dilution for immunofluorescence.

**Cell Culture**

Cells used in all experiments were monomorphic procyclic form *T. brucei* Lister strain 427 or a tetracycline-inducible strain 427-based strain generated by introducing the Single Marker Oxford plasmid (30). 427-based cell lines were grown at 27 °C in Beck's Medium (Hyclone, Logan, Utah) supplemented with 500 µg/mL penicillin-streptomycin-glutamine (Hyclone), 10% fetal bovine serum (Gemini Bioproducts, West Sacramento, CA), and 10 µg/mL gentamycin (ThermoFisher Scientific, Waltham, MA). SmOx-based cell lines were cultured at 27 °C in Beck's Medium supplemented with 500 µg/mL penicillin-streptomycin-glutamine, 1 µg/mL puromycin, 10% tetracycline-free fetal bovine serum (RnD Systems, Minneapolis, MN), and 10 µg/mL gentamycin. Cell counts were determined using a Z2 Coulter Counter particle counter (Beckman Coulter, Brea, CA).

**Cloning and Cell Line Assembly**

All DNA constructs were assembled using PCR amplified inserts followed by restriction-ligation or Gibson Assembly into vectors as previously described (1). The mClv3 endogenous tagging construct for PFR2 (Tb927.8.4970) was targeted to the endogenous loci using the last 500 bp of the coding region of PFR2 and the first 500 bp of the 3' untranslated region. Sequences suitable for RNAi were obtained using the RNAit software (https://dag.compbio.dundee.ac.uk/RNAit/) and inserted into a modified pLEW100 vector

(1, 2). 400-600 bp of the coding sequence were targeted for each of the following proteins: PFR2-mClv3: bp 249-671 of mClv3 sequence; TOEFAZ1 (Tb927.11.15800): bp 578-1090. TbPLKas cells used were previously generated (3).

Constructs were transfected into cells using an electroporator (GenePulser xCell, Bio-Rad, Hercules, CA). Clonal cell lines were isolated by limiting dilution and selection with the appropriate resistance marker. All cell lines were verified using loci PCR, immunofluorescence (IF), and western blotting.

For TbPLKas experiments, a cell line ectopically expressing the GFP variant mClv3 in the cytosol was generated by inserting mClv3 coding sequence into a pXS2 plasmid and inserting into 427-derived cells to form 427 sentinels. For the TOEFAZ1 RNAi experiments, a cell line with an inducible cytosolic mNeonGreen was generated by inserting mNeonGreen coding sequence into a pLew100 vector and inserting into SmOx cells to form inducible SmOx sentinels.

**Immunofluorescence**

Cells were harvested by centrifugation at 2400 × g for 5 min and washed once in PBS. Cells were centrifuged onto coverslips and fixed in prechilled methanol at -20 °C for 20 min. Coverslips were then washed 3 times in PBS, followed by blocking buffer (5% goat serum (Gibco, ThermoFisher Scientific, Waltham, MA) in PBS. Coverslips were incubated in primary antibody diluted into blocking buffer for 1 h at room temperature (RT), after which cells were washed three times in PBS. Coverslips were then incubated with secondary antibody conjugated to Alexa Fluor−568 (Life Technologies, Carlsbad, CA) for 1 h at RT. After 3 washes in PBS, coverslips were mounted in Fluoromount G with DAPI (Southern Biotech, Birmingham, AL).

**Microfabrication**

We used a previously described approach (4–7)to design squared microchambers to house *T. brucei*. The squared microchambers (more than 10,000 per array) were designed in AutoCAD (Autodesk, San Rafael, CA) and CleWin (Delta Mask). Briefly, silicon dioxide wafers were cleaned using an adapted RCA cleaning procedure in 1:1:5 $H_2O_2$:$NH_4OH$:$H_2O$. A 1:10 hexadimethylsiloxane:isopropanol solution was incubated on top of the clean wafer for 10 min. The solution was then removed by spincoating and the wafer was heated to 95°C on a heating plate to remove excess solvent and to activate HDMS for adhesion of the photoresist. We spincoated positive Shipley photoresist 1828 (Microchem) either once or twice to achieve appropriate heights ranging from 2-6µm onto the activated silicon wafer. The photoresists were processed according to the manufacturer's instructions. The microchamber design was then directly written onto the photoresists using a µMLA tabletop maskless aligner (Heidelberg Instruments). After exposure the photoresist was developed with a drop of MF-321 (MicroChem) for ~2 min on top of the array, followed by blow drying under a stream of nitrogen. The master was silanized overnight under vacuum with a drop of heptadecafluoro-1,1,2,2-tetrahydrodecyl trichlorosilane (Gelest Inc.). The array of microchambers was then replicated into PDMS (Sylgard184) using a ratio of 10:1 base:curing agent. The PDMS stamp was cured overnight under vacuum at 65 °C. The PDMS mold was then peeled off from the silicon master and can be used repetitively to create the pattern in medium-agarose.

#### RNAi

RNAi was induced in cell lines containing pTrypSon-based lhRNAi plasmids (49). For live-cell experiments, cells were seeded at $2\times10^6$ cells/mL and induced with 1 µg/mL doxycycline (ThermoFisher Scientific). For PFR2-mClv3 experiments, RNAi against mClv3 was induced for 12 h before plating. TOEFAZ1 RNAi and pLew100-mClv3 sentinels were induced for 16 h before plating. Cells were harvested by centrifugation at 800 × g for 10 min RT. Cells were resuspended in 1 mL of base SmOx media with 1 µg/mL doxycycline and 100 µM reduced L-glutathione.

Cell growth was monitored for PFR2-mClv3 RNAi and inducible pLew100-mClv3 cells to ensure no growth defects were present. Cells were grown in either 1 µg/mL doxycycline, or with an equal volume of 70% ethanol as a vehicle control. Cell growth was monitored every 24 h by counting. Every 48 h, cultures were reseeded in fresh media with either doxycycline or 70 % ethanol. Generation plots for the TOEFAZ1 RNAi and inducible mClv3 expression depict averages of triplicate experiments, with standard deviation error bars shown.

#### Statistical Analysis

Graphs with corresponding statistical analysis in figure legends were generated using GraphPad Prism 9 software version 9.1.1 (GraphPad Software). Unpaired two-tailed students *t* tests were used to determine if differences seen between groups were statistically significant at the $p < 0.05$ significance level. A one-sided Fisher exact test was performed for all PFR2-mClv3 RNAi, TbPLKas and TOEFAZ1 RNAi experiments to ensure enough replicates were performed to provide sufficient statistical power to detect a significant difference at a $p<0.05$ significance level.

#### Supplementary Figure Legends/List

##### Figure S1. Assembly of the live-cell imaging apparatus.

**A.** Representative image of a silicon wafer generated through photolithography. The wafer is used to generate a PDMS stamp that contains the inverse of the well sizes. **B.** Agarose (denoted by arrow) is overlaid on the PDMS stamp to generate the microwells. **C.** Cells are plated into the Lab-Tek chamber and the cut-to-size agarose microwells (denoted by arrow) are overlaid on the cells. **D.** Grid is weighed down and sealed with mineral oil to prevent evaporation. A glass slide is adhered with high vacuum grease (Dow Corning) to seal the chamber slide. **E.** Representative DIC image of SmOx cells in 100×100×5 μm wells, imaged with a 20×/0.8 NA air lens.

##### Figure S2 PFR2-mClv3 RNAi L8C4 colabel and growth curve

RNAi against the single PFR2-mClv3 allele does not affect cell growth or the paraflagellar rod. **A.** RNAi against PFR2-mClv3 was induced for 12 h. Cells were harvested and methanol fixed and stained using L8C4 antibody to label the PFR. Native fluorescence of PFR2-mClv3 is shown. Depletion of the PFR2-mClv3 allele through RNAi results in a normal new PFR structure with no PFR2-mClv3 signal. **B**. RNAi against PFR2-mClv3 was induced for 4 days with 1 μg/ml doxycycline or 70% ethanol as a vehicle control. Cell concentration was counted every 24 h. Graph depicts three independent experiments; error bars depict standard deviation.

**Movie S1 24 h time course captures multiple rounds of cell division**

SmOx cells were plated in 100×100×5 μm wells and imaged with a 40×/1.3 NA oil lens. DIC images were captured every 10 minutes for 24 h. Video depicts images shown in Figure 1A. Video presented at 5 frames per second.

**Movie S2 Higher sampling rate captures cell cycle events with high resolution**

SmOx cells were plated in 50×50×5 μm wells and imaged with a 40×/1.3 NA oil lens. DIC images were captured every 15 s for 2 h. Video depicts images shown in Figure 1B. Video is presented at 5 frames per second.

**Movie S3 SmOx Differential Division**

SmOx cells were plated in 100×100×5 μm wells and imaged with a 20×/0.8 NA lens. DIC images were captured every 10 min for 24 h. Video depicts images shown in Figure 2. Video is presented at 5 frames per second.

**Movie S4 PFR2-mClv3 RNAi results in differentially labelled daughter cells**

PFR2-mClv3 cells were induced for mClv3 RNAi for 12 h and were plated in 100×100×5 μm wells. Split-view system allows for simultaneous capture of DIC and FITC images. Images were taken with a 40×/1.3 NA oil lens every 10 min for 24 h using 30 ms exposure and 4% LED power with 2×2 pixel binning. Video depicts timelapse shown in Figure 3A. Video is presented at 5 frames per second.

**Movie S5 Small molecule inhibition of TbPLKas with 427 Sentinels**

TbPLKas and 427 mNeonGreen-Sentinels were incubated with 2.5 μM 3MB-PPi for 3 h before plating in 100×100×5 μm wells. Split-view system allows for simultaneous capture of DIC and FITC images. Images were taken with a 40×/1.3 NA oil lens every 10 min for 24 h using 30 ms exposure and 4% LED power with 2×2 pixel binning. Video depicts time lapse shown in Figure 4A. Video is presented at 5 frames per second.

**Movie S6 TbPLKas inhibition results in a looped new flagellum**

TbPLKas cells were incubated with 2.5 μM 3MB-PPi for 3 h before plating in 100x100x4 μm wells. DIC images were taken every 5 min for 24 h with a 40×/1.3 NA oil lens using 30 ms exposures. Video depicts time lapse shown in Figure 4D. Videos are presented at 5 frames per second.

**Movie S7 TOEFAZ1 RNAi produces an unproductive posterior furrow**

TOEFAZ1-RNAi was induced for 16 h prior to plating in 50×50×5 μm wells. DIC images were taken every 5 min for 20 h with a 40×/1.3 NA oil lens with 30 ms exposures. Video depicts time lapse shown in Figure 5. Video is presented at 5 frames per second.

**Movie S8 TOEFAZ1 RNAi and SmOx Sentinel**

TOEFAZ1-RNAi and SmOx mClv3-sentinel cells were induced for 16 h prior to plating in 100×100×5 µm wells. Split-view system allows for simultaneous capture of DIC and FITC images. Images were taken every 10 min for 20 h with a 40×/1.3 NA oil lens using 30 ms exposure and 30% LED power. Videos show time lapse depicted in Figure 6. Video is presented at 5 frames per second.

**SUPPLEMENTAL REFERENCES**

1. McAllaster MR, Sinclair-Davis AN, Hilton NA, Graffenried CL de. 2016. A unified approach towards Trypanosoma brucei functional genomics using Gibson assembly. Mol Biochem Parasit 210:13 21.

2. Redmond S, Vadivelu J, Field MC. 2003. RNAit: an automated web-based tool for the selection of RNAi targets in Trypanosoma brucei. Mol Biochem Parasit 128:115 118.

3. Lozano-Núñez A, Ikeda KN, Sauer T, Graffenried CL de. 2013. An analogue-sensitive approach identifies basal body rotation and flagellum attachment zone elongation as key functions of PLK in Trypanosoma brucei. Mol Biol Cell 24:1321 1333.

4. Renner LD, Weibel DB. 2011. Cardiolipin microdomains localize to negatively curved regions of Escherichia coli membranes. Proc National Acad Sci 108:6264–6269.

5. Eun Y-J, Ho P-Y, Kim M, LaRussa S, Robert L, Renner LD, Schmid A, Garner E, Amir A. 2018. Archaeal cells share common size control with bacteria despite noisier growth and division. Nat Microbiol 3:1 9.

6. Wong F, Renner LD, Özbaykal G, Paulose J, Weibel DB, Teeffelen S van, Amir A. 2017. Mechanical strain sensing implicated in cell shape recovery in Escherichia coli. Nat Microbiol 2:17115.

7. Renner LD. 2019. Engineering Bacterial Shape Using Soft Matter Microchambers. Curr Protoc Chem Biology 11:e59.

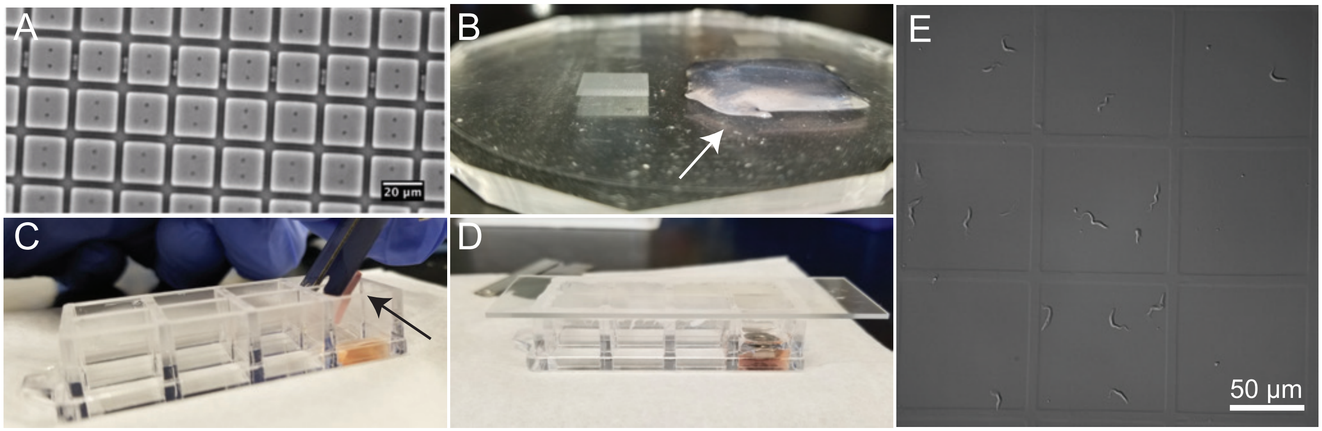


**Supplemental Figure 1**

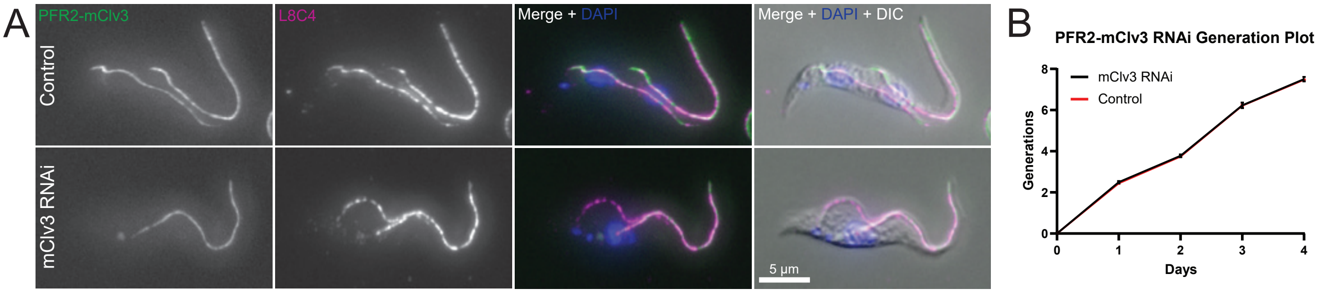


### Supplemental Figure 2
